# Peptide-aware chemical language model successfully predicts membrane diffusion of cyclic peptides

**DOI:** 10.1101/2024.08.09.607221

**Authors:** Aaron L. Feller, Claus O. Wilke

## Abstract

Language modeling applied to biological data has significantly advanced the prediction of membrane penetration for small molecule drugs and natural peptides. However, accurately predicting membrane diffusion for peptides with pharmacologically relevant modifications remains a substantial challenge. Here, we introduce PeptideCLM, a peptide-focused chemical language model capable of encoding peptides with chemical modifications, unnatural or non-canonical amino acids, and cyclizations. We assess this model by predicting membrane diffusion of cyclic peptides, demonstrating greater predictive power than existing chemical language models. Our model is versatile and can be extended beyond membrane diffusion predictions to other target values. Its advantages include the ability to model macromolecules using chemical string notation, a largely unexplored domain, and a simple, flexible architecture that allows for adaptation to any peptide or other macromolecule dataset.

## 1 Introduction

Therapeutic peptides have gained attention as promising clinical agents due to their diverse chemical properties and their efficacy in treating a variety of diseases.^1–3^ Advances in peptide production and modification technologies have facilitated the development of drug targeting strategies involving cyclization reactions and the alteration of both side-chain and backbone chemistries beyond those found in nature. ^4–11^ Existing deep-learning models have been successful in modeling small molecule drugs and large proteins.^12,13^ Furthermore, protein language models have been finetuned for inference tasks on natural peptides. ^14^ However, these frameworks fall short in peptide drug discovery applications due to limited diversity in pretraining data and restrictions imposed by natural amino acid vocabularies. The increasing diversity in synthetic peptide chemistries necessitates robust computational methodologies capable of accurately encoding and predicting their properties.

Despite an interest in developing peptide drugs for intracellular targets ^15,16^ the landscape of available models is sparse. While models for predicting membrane penetration of small-molecule drugs and natural, linear peptides are abundant and have achieved high accuracy,^7,17,18^ there are no pretrained language models for modeling of cyclic or modified peptides. The predictive modeling of these complex molecules has had to rely on molecular dynamics simulations, which are computationally expensive.^19^ Moreover, Lipinski’s rule of five that serves as reliable guidelines for membrane diffusion of small molecules fails to generalize to peptides. ^20,21^ Therefore, a new modeling approach is required for peptide-based drug discovery, one that can encode diverse modifications, unnatural amino acids, and cyclizations.

Prior modeling of membrane permeation of cyclic peptides with deep learning models has resulted in promising results using various combinations of graph neural networks, transformer models, and molecular descriptors.^22–24^ However, in these examples the models were not pretrained, and the training and test sets were randomly selected with a large overlap in chemical space. Both of these choices can lead to an overfit model that can fail to accurately predict membrane permeation on molecules that fall out of distribution for the training set. This is apparent from a marked decrease in performance on some independent test sets.^23^

Here, we present PeptideCLM, a chemical language model trained on both small molecules and peptides. This model was designed with the aim of generalizing to unseen data by first pretraining a chemical language model using a masked language modeling approach with an atomic string notation language known as Simplified Molecular-Input Line-Entry System (SMILES). A pretraining database of 23M molecules, including 10M peptides with chemical modifications, non-canonical amino acids, and cyclic structures, was created to increase training data to a level that is consistent with language modeling scaling laws. We assess generalizability by predicting permeability on peptides clustered using k-means to select training, validation, and test sets. On this dataset, our model achieves higher predictive accuracy for membrane diffusion than existing chemical language models. PeptideCLM has been released on Hugging Face for simple implementation, providing a versatile tool for peptide research and drug discovery.

## 2 Methods

### Pretraining dataset generation

Two distinct datasets were curated for chemical language model (CLM) pretraining. The first consists of 10 million small molecules from Pub-Chem,^25^ as released with the ChemBERTa model,^26^ and 2.2 million small molecules from SureChEMBL,^27^ filtered for single-molecule entries. The 10M PubChem dataset was selected as it was previously used to successfully pretrain a chemical language model. We expanded this data with 2.2M molecules from SureChEMBL in an effort to infuse the model with knowledge of medically relevant chemical structures, as all molecules in this dataset were taken from the patent literature. The second dataset consists of 825,632 peptides from SmProt^28^ and 10 million generated non-natural peptides. SmProt is a database of peptides (*<*100 amino acids) annotated from ribosome profiling across 8 species, and is the largest available database of natural small proteins and peptides. We then added 10 million generated modified peptides (Figure 1) using an updated version of CycloPs.^29^ This data was included to provide the model with examples of non-natural peptides and increase training data to near that of the small molecule dataset.

**Figure 1:**
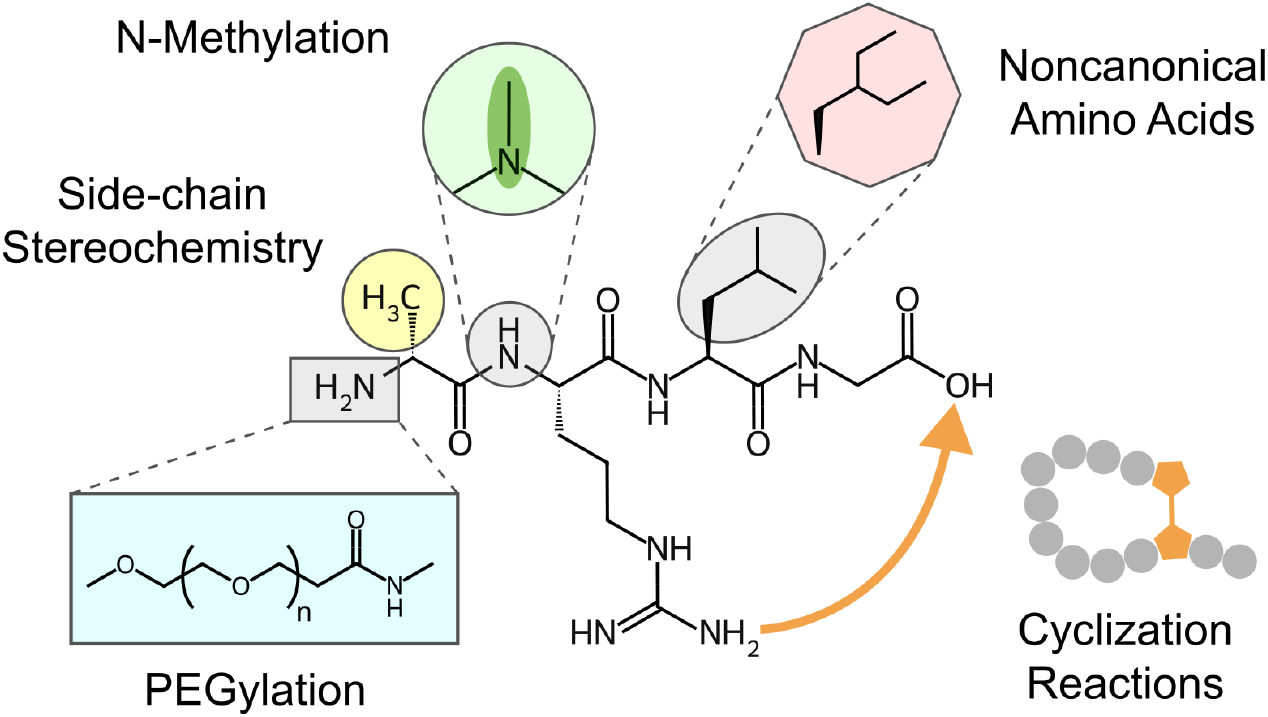
Example modifications of the tetrapeptide Ala-Arg-Leu-Gly. The peptide structure is depicted with five modifications used for synthetic data generation: (clockwise) N-methylation of a backbone amine, substitution of alanine for the synthetic amino acid diethylalanine, cyclization of asparagine to the carboxyl terminus, PEGylation of an amino group, and altered side-chain sterochemistry.

All non-natural peptides were generated de novo using stochastic sampling. First, we expanded the capabilities of the CycloPs software by adding 100 types of non-natural amino acids from SwissSidechain. ^30^ We then uniformly sampled amino acids to generate peptides of 100 or fewer amino acids consisting of 90% natural and 10% unnatural amino acids. Furthermore, the stereochemistry at 10% of the *α* carbons was changed to dextro. Finally, all generated peptides were assigned a cyclization reaction, randomly sampling from head-to-tail, sidechain-to-sidechain, sidechain-to-head, sidechain-to-tail, or a disulfide bridge. We used CycloPs to convert the amino acid notation to chemical strings using Simplified Molecular-Input Line-Entry System (SMILES) notation.^31^ If cyclization was not possible given the available amino acid residues, the molecule was left linear. Once the chemical strings were generated, they were modified to add N-methylation on a random 20% of backbone amines on 20% of peptides (4% total) followed by attachment of polyethylene glycol (PEGylation) to 20% of the peptides, with monomer lengths between 1-4 attached at a random free amine.

### Tokenization scheme

A custom tokenization method, derived from SMILES Pair Encoding,^32^ was developed to optimally represent peptide chemistries from SMILES strings (Figure 2A). A pretokenizer first identified multi-character tokens such as bromine and chlorine as Br and Cl, which are distinct from boron and carbon represented by B and C. Subsequently, token assignment of chemical motifs was performed through n-gram analysis of up to 5 characters over the 10 million PubChem SMILES strings. This process identified a total of 581 unique tokens, resulting in 586 tokens total when including the 5 special tokens [PAD], [UNK], [CLS], [SEP], and [MASK] which represent end-sequence padding, unknown character, beginning of sequence, end of sequence, and a masked position, respectively. Among the 581 tokens, 53 are single-character tokens, 150 are multi-character tokens (e.g., CCC, Clc, occ), 124 are bracketed ions (e.g., [Ag^+^], [Ag], [Al^*−*3^]), and 254 are parenthetical motifs (e.g., (/CCl), (=C), (CCCN)).

**Figure 2:**
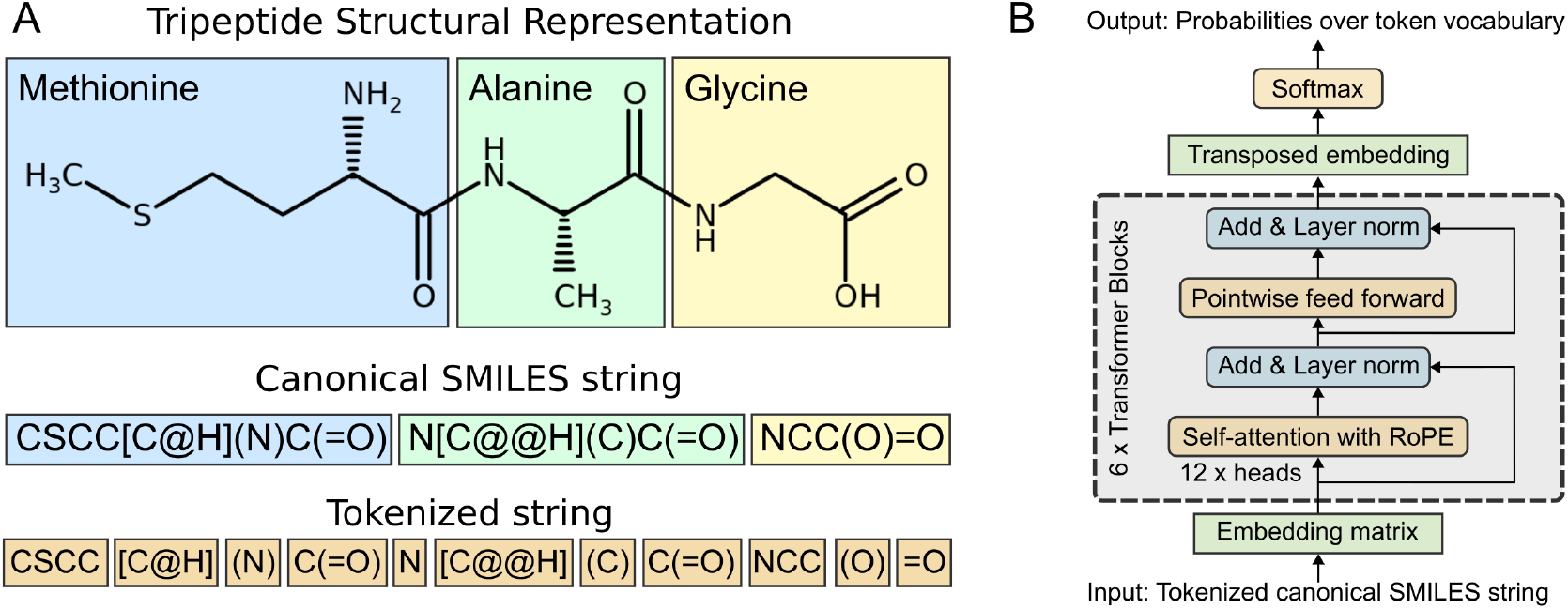
Schema for tokenization of a tripeptide and implementation of a BERT-style architecture for encoding peptides. (A) 2D-structure representation colored by amino acid. SMILES canonical representation generated using RDKit with same coloring. Representative tokens for input into a PeptideCLM. (B) The achitecture for the RoFormer BERT model used as the framework for PeptideCLM. The model contains six transformer blocks with twelve attention heads per block, post-layer normalization, and a standard language modeling head with a softmax distribution over the vocabulary index for token prediction.

### Model architecture and pretraining

PeptideCLM is a BERT-style transformer architecture with Rotary Position Embedding (RoPE) based on the RoFormer implementation described by Su (2024).^33^ Our implementation of RoFormer employs 12 attention heads and 6 layers (Figure 2B), with a hidden embedding dimension of 768 and an intermediate feedforward layer size of 3072, matching the configurations used in GPT-3^34^ and BERT.^35^

With this architecture, three models were trained to assess the impact of pretraining data. A “full” model was trained for 10 epochs on the combined dataset of small molecules and peptides (Figure 3A). Two additional models were trained for 20 epochs each on either the peptide portion or the small molecule portion of the pretraining data. These ablation models were trained for 20 epochs to reach a similar number of training steps as used in the full dataset. For all models, training was conducted using a masked prediction task, wherein 15% of the sites were masked. Each sequence underwent an 80/10/10 split for masking, corrupting with a random token, or leaving the sequence unmasked, respectively, as described in the BERT methodology.^35^ The hyperparameters for pretraining PeptideCLM were a batch size of 64 and a learning rate of 5 *×* 10^*−*5^, chosen based on preliminary experiments indicating these values provided a good balance between convergence speed and stability. The holdout dataset for pretraining validation comprised 0.5% of the full model and 1% of the chemical and peptide models, amounting to roughly 115k SMILES strings per model. Validation was performed at intervals corresponding to every 20% of an epoch to ensure consistent model performance and early detection of potential overfitting. The final checkpoint was used for downstream application. Full hyperparameters for training are reported in Supplemental Table 3.

**Figure 3:**
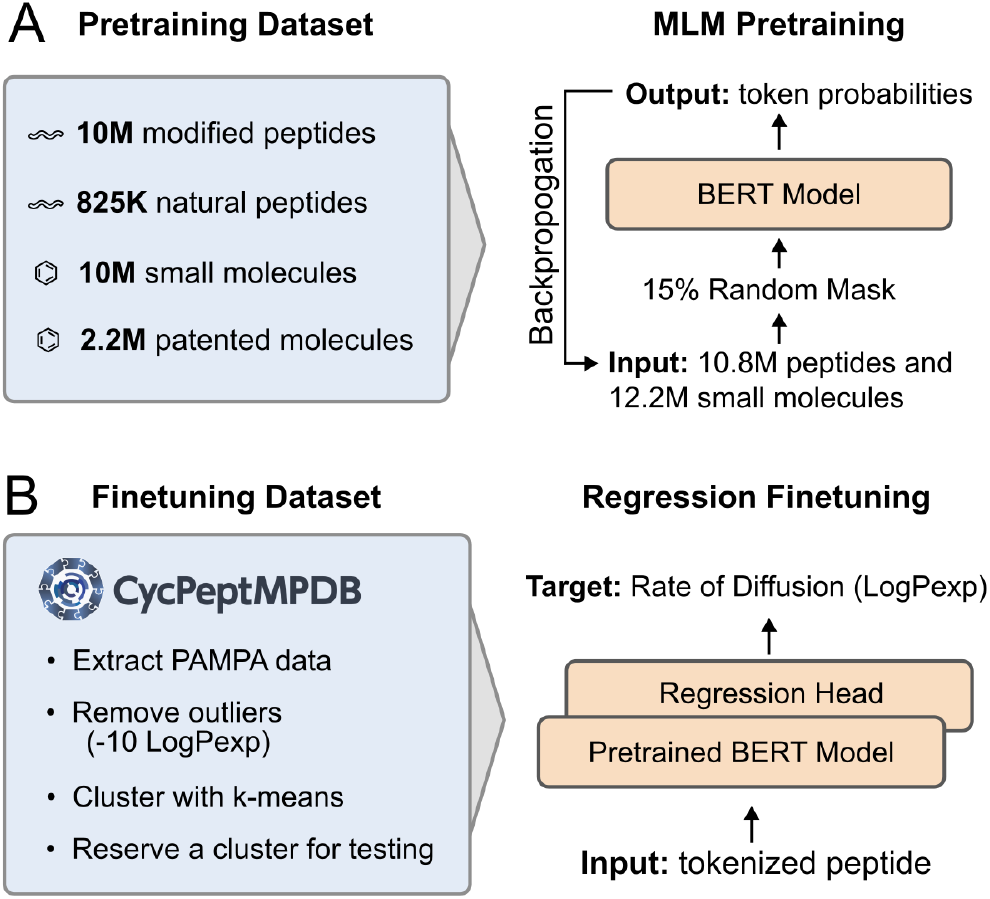
Overview of pretraining and finetuning workflows. (A) Pretraining Dataset and MLM Pretraining: The dataset includes 10M modified peptides, 825K natural peptides, 10M small molecules, and 2.2M patented molecules. The BERT model is pretrained with a masked language model (MLM) objective. (B) Finetuning Dataset and Regression Finetuning: PAMPA data from CycPeptMPDB is clustered, with outliers removed and one cluster reserved for testing. The pretrained BERT model is finetuned with a regression head to predict diffusion rate (Log*P*_*exp*_).

### Finetuning data cleaning

The Cyclic Peptide Membrane Permeability Database (CycPeptMPDB)^36^ contains measured permeability of nearly 7,500 cyclic peptides. The data is curated from 45 peer-reviewed publications and 2 pharmaceutical company patents, each contributing valuable data on cyclic peptides. A significant portion of the studies within CycPeptMPDB include Parallel Artificial Membrane Permeability Assay (PAMPA) data, providing insights into the passive permeability of cyclic peptides. Many of these measurements were performed by independent researchers, adding a diverse range of experimental conditions and methodologies.

Most permeability measurements in CycPeptMPDB were conducted using Liquid Chromatography-Mass Spectrometry (LC/MS), known for its high sensitivity and accuracy in detecting trace amounts of compounds. In a few cases, Liquid Chromatography-Ultraviolet Detection (LC/UV) was used. Each dataset entry is cited in CycPeptMPDB, linking back to the original publications, ensuring transparency and enabling users to trace the source of each measurement. We have relied on this data as it is reported in the combined database, though general error rates for LC/MS are 1–5% relative standard deviation and are higher for LC/UV in the range of 5–10% relative standard deviation.

For our analysis, we subset CycPeptMPDB to only molecules that had results from PAMPA, as the full database also includes results from cell-based permeability assays including colorectal adenocarcinoma cells (Caco2), Madin-Darby Canine Kidney cells (MDCK), and Ralph Russ Canine Kidney cells (RRCK). All data points that were categorized as “undetectable” and marked as *−*10 in CycPeptMPDB were removed due to the potential of aggregation or some other data collection error, which may lead to PAMPA results differing from the true permeability of the peptide. The distribution of PAMPA scores for all selected peptides is shown in Figure S1.

### Train/test clustering

In an effort to generate training and test sets that are distinct in chemical space, we used our pretrained model to conduct k-means clustering and used a *leave-one-cluster-out* approach for train-test splits with k-fold validation. We first generated embeddings of all peptides with PAMPA scores using the pretrained PeptideCLM. We reduced the dimensions of these embeddings using principal component analysis to a minimum number that retained 99% of the variance, a total of 153 components. We clustered the peptides into train/validation/test using k-means clustering on the reduced embedding space. We chose to use k-means as the method and a total of six clusters based on high silhouette score^37^ (Figure S2), low Davies-Bouldin Index^38^ (Figure S3), and high Calinski-Harabasz Index^39^ (Figure S4). We treated each cluster as an independent holdout set, iterating through all six clusters. For each test cluster, five models were trained on the remaining training clusters, using 5-fold cross-validaiton to predict permeability score (Log*P*_*exp*_). Training was allowed to proceed for a maximum of 10,000 steps and the final weights were taken from the checkpoint exhibiting the lowest mean squared error (MSE) on the validation cluster.

### Model finetuning

For each of the 11 models we assessed, we attached a fully connected feed-forward layer—matching the width of the model’s hidden state—to replace the language modeling head (Figure 3B), predicting a single output value. The entire network was finetuned on each model using mean squared error (MSE) as the loss function. We predicted the permeability score on the holdout test cluster using the mean prediction from the k-fold models. This training–prediction method was repeated six times, with each cluster used as the holdout test data one time.

For model scoring, we treated the predictions as a binary classification task by setting a cutoff at 10^*−*5.5^ Log*P*_*exp*_. Peptides with Log*P*_*exp*_ values below this threshold were classified as non-permeable and those at or above it were classified as permeable. This value was selected as it corresponded to a marked increase in oral bioavailability of drug-like compounds in Veber, et al.^40^ This cutoff also provides a more even split between permeable and non-permeable than the value of 10^*−*6^ that was proposed in CycPeptMPDB.^36^ We then generated Receiver Operating Characteristic (ROC) curves and used the Area Under the Curve (AUC) as the scoring metric. Additionally, we used the AUC on Precision-Recall (PR) curves and calculated the root mean squared error (RMSE) to further evaluate model performance. The mean and standard deviation for these metrics across six holdout test clusters were calculated for each model.

Finetuning of all models on our subset of the CycPeptMPDB data was performed with the following hyperparameters: a learning rate of 5 *×* 10^*−*6^, a dropout rate of 0.15, and a weight decay of 0.001. The batch size was set to 16, and training was conducted for a maximum of 10,000 steps. Testing on the validation cluster was performed after each epoch to monitor performance. Full hyperparameters can be found in Table S3.

### Framework and hardware

All model training and inference were conducted on a system running Linux 20.04.6 LTS equipped with 8x AMD Navi 10 (Radeon RX 5700 XT) GPUs. Pretraining was conducted on 8 GPUs using distributed data parallelism. Finetuning was conducted in parallel with multiple models/folds running on a single GPU. The machine learning framework employed for these tasks was PyTorch, ^41^ utilizing PyTorch Lightning^42^ to manage the training process.

## 3 Results

Three models were pretrained to evaluate the impact of the synthetic peptide data on pretraining loss and the downstream task of predicting membrane diffusion. Results from the pretraining task showed that training the model with small molecules from Pubchem and SureChEMBL had a steeper descent for lower cross-entropy loss (CEL) and higher maximum accuracy (Figure 4). The peptide-only model achieved the slowest loss descent and poorest accuracy. Pretraining on a combined dataset of peptides and small molecules resulted in a CEL descent that was intermediate between the two individual models, achieving a final accuracy comparable to the peptide-only model. This result is expected, as randomization of peptides results in tokens that are equally likely. For example, the chirality of the *α*-carbon cannot be learned, as the chirality was random and could not be inferred from the rest of the sequence.

**Figure 4:**
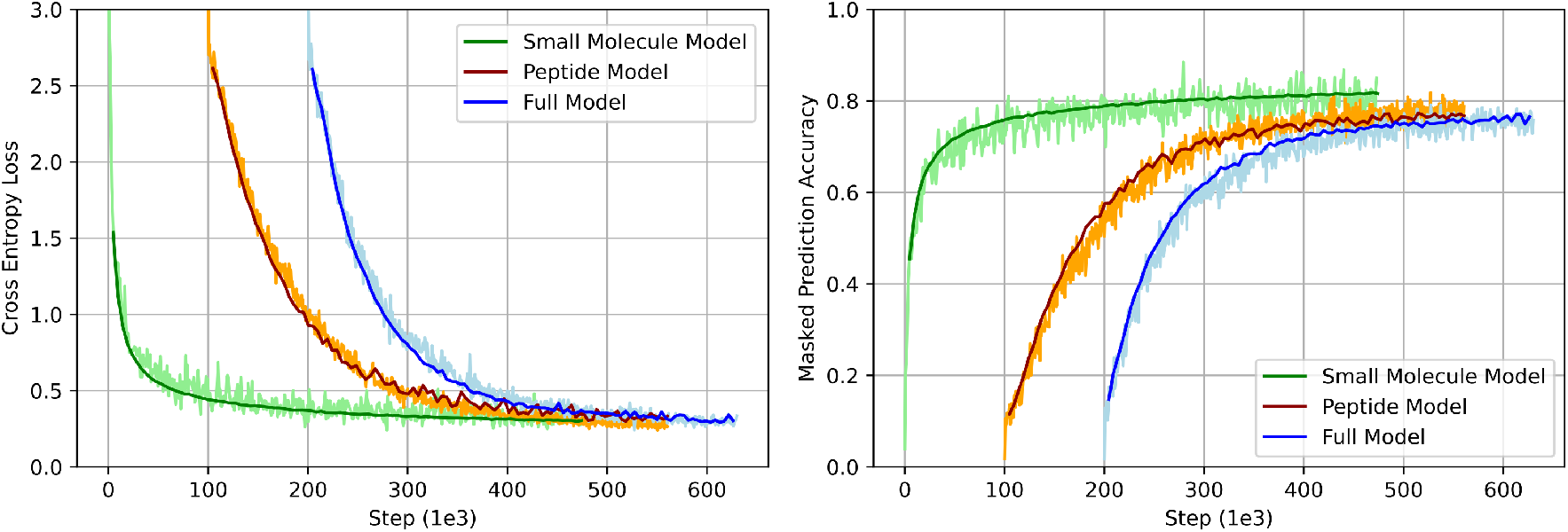
Cross-Entropy Loss and Macro-Averaged Accuracy in Training a BERT-Style Model with Masked Prediction. Cross-entropy loss (left) is computed between the softmax of the token probabilities and the one-hot encoded original sequence. The loss is determined as the sum of the logarithmic loss *−*(*y* log(*p*) + (1 *− y*) log(1 *− p*)) across all masked positions. Macro-averaged accuracy (right) is calculated as the average accuracy, averaged across classes. The validation set comprised a randomly selected 0.5% of the full dataset, or 1.0% of the peptide or small molecule datasets. Peptide and Full models are shifted right 100*e*3 and 200*e*3 steps, respectively.

The performance of PeptideCLM, ChemBERTa-2, and ChemBERTa was evaluated on their ability to predict the membrane diffusion of cyclic peptides. First, the data was split into six clusters (Figure 5A) by applying k-means clustering on the top 153 principal components of the pretrained PeptideCLM peptide embeddings. The models were then finetuned to predict the logarithm of the experimental partition coefficient (Log*P*_*exp*_) in a regression manner. This was performed across 5-folds of the data for a maximum of 10,000 steps, with checkpointing at the end of each epoch. The final checkpoint was selected based on the lowest MSE observed on the validation set. The models were then used to classify membrane penetration by setting a cutoff for Log*P*_*exp*_ of *−*5.5 (1.0*×*10^*−*5.5^ cm/s) or higher for membrane penetrating peptides, and lower for non-penetrating. Final test metrics were calculated as the mean pooled prediction for the five models.

**Figure 5:**
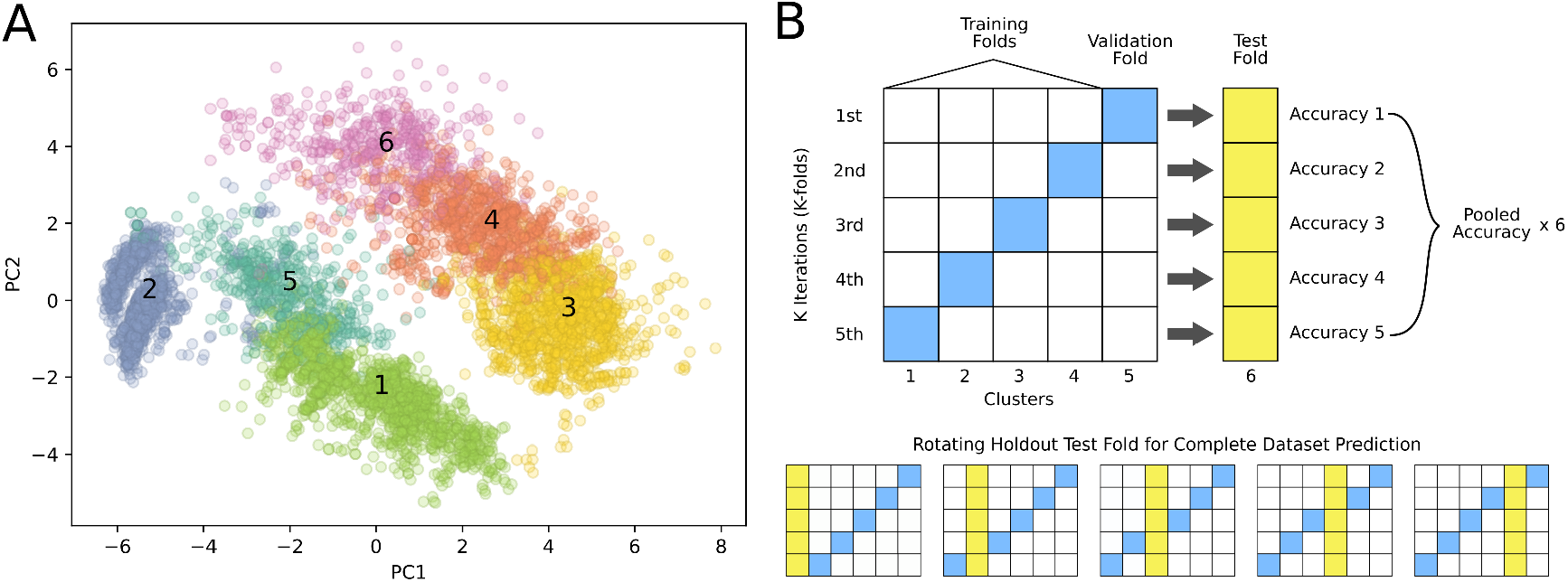
Clustering of CycPeptMPDB peptide embeddings and *leave-one-cluster-out* validation method. Principal Components Analysis (PCA) was applied to peptide embeddings generated by the pretrained PeptideCLM model, followed by k-means clustering to identify six distinct clusters. (A) The PCA plot visualizes the distribution of peptides across the six clusters. (B) One cluster (yellow) is held out as the test set, while the remaining clusters are used for training (white) and validation (blue). This rotation continues through all six clusters, ensuring that each cluster is tested once while the others are used for training. For each iteration, the accuracy is calculated for the test set (yellow) with the average prediction for all holdout sets used as the model score.

The entire finetuning process was repeated six times, rotating through all six clusters as the holdout test set (Figure 5B). The mean Receiver Operating Characteristic (ROC) and Precision-Recall (PR) curves for each holdout set were generated and area under the curve (AUC) was used for quantification (Figure 6). We also calculated the root mean squared error (RMSE) between the mean prediction for each holdout test set. The mean and standard deviation for each model is reported in Table 1. PeptideCLM performed best with a ROC-AUC of 0.781 and a PR-AUC of 0.738. All ROC curves for PeptideCLM are presented in Supplemental Figures S5-10.

**Table 1:**
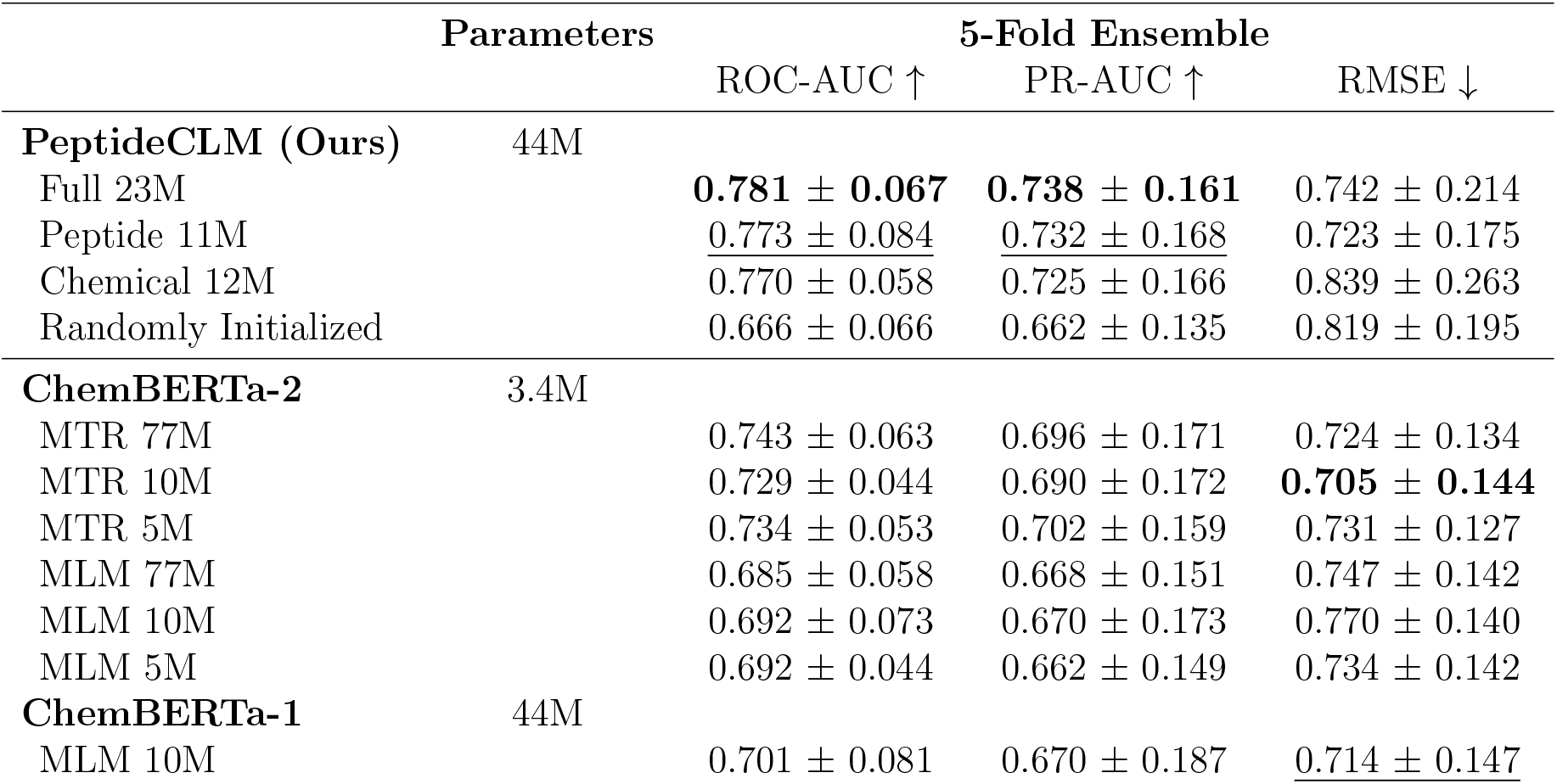
Performance comparison of PeptideCLM and ChemBERTa models. The highest-scoring model is typeset in bold, and the second-highest scoring model is underlined. The models evaluated include PeptideCLM and ChemBERTa variants trained with different pretraining datasets. PeptideCLM variants are noted by their pretraining dataset; Full 23M was trained on the combined data of both 11M peptides and 12M small molecules. The ChemBERTa-2 models are differentiated by their pretraining dataset size, where 77M, 10M, and 5M refer to the number of small molecules in millions used during pretraining. MTR stands for Multi-Task Regression and MLM stands for Masked Language Modeling. RMSE indicates the root mean squared error of predictions.

**Figure 6:**
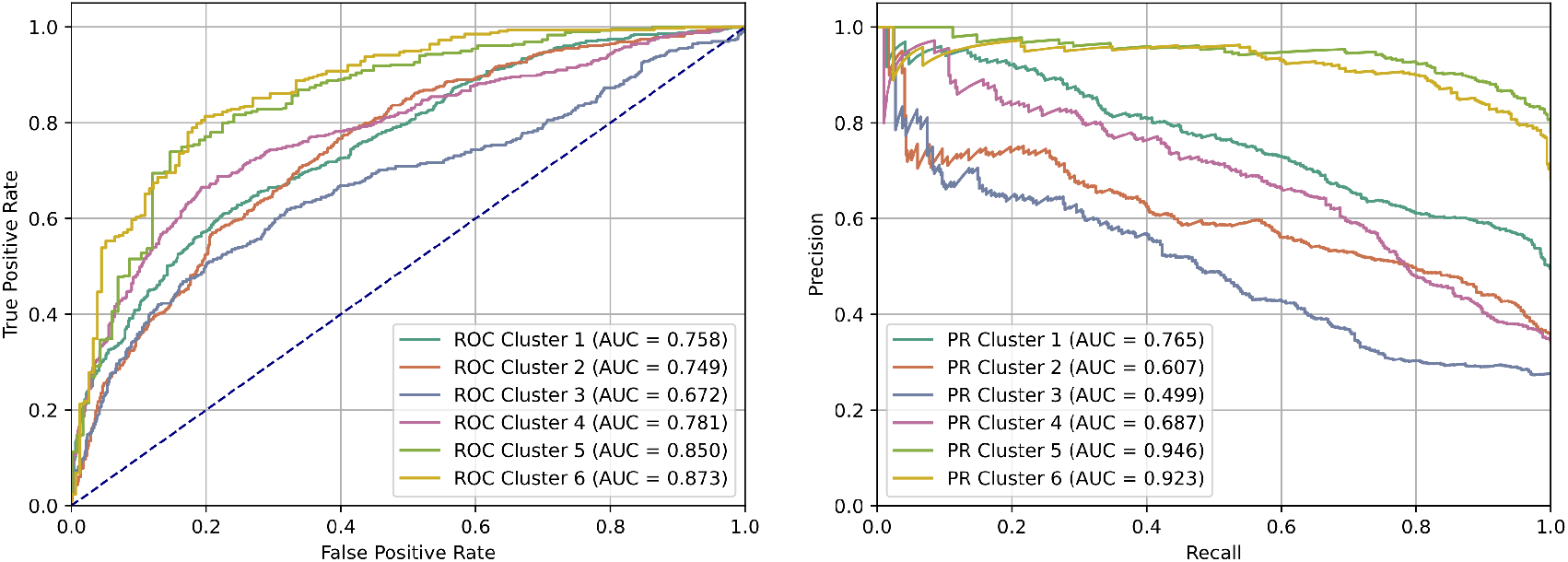
Receiver-operator curves and precision-recall curves of finetuned PeptideCLM for predicting membrane penetration. Receiver-operator curves (Left) and precision-recall curves (right) of the finetuned PeptideCLM model using the mean of five models generated during cross-validation. Each curve represents one evaluation on a holdout test cluster and is generated by taking the mean pooled prediction of the five-fold validations.

Our model outperformed the other available models, regardless of the pretraining data used. We observed an increase in ROC-AUC with the addition of peptide data, suggesting that in-domain pretraining data can enhance downstream finetuning and prediction. The second-highest scoring model was ChemBERTa-2 MTR across all dataset sizes, followed by the original ChemBERTa model in third place. The ChemBERTa-2 MLM models performed the worst. It is notable that the pretraining dataset size did not correspond directly to an improvement on our finetuning task, as shown with the ChemBERTa-2 models.

## 4 Discussion

In this study, we have developed a peptide-specific chemical language model, PeptideCLM, capable of encoding peptides with various chemical modifications, including non-canonical amino acids and cyclic structures. Our model has demonstrated superior performance in predicting membrane diffusion from SMILES strings of an embedding-based holdout set of cyclic peptides, outperforming the existing chemical language models capable of performing this task, namely ChemBERTa and ChemBERTa-2. These results highlight the potential of this modeling approach in addressing the non-trivial challenges in peptide modeling. In particular, we have highlighted the promise of encoding complex macromolecules using a chemical string notation.

Our results indicate that a chemical language model (CLM), when pretrained on a diverse dataset of natural and synthetic peptides, exhibits an enhanced ability to generalize to out-of-distribution peptides. The higher evaluation scores demonstrate the model’s robustness in predicting membrane permeability on out-of-distribution targets, which is crucial for designing peptides to interact with intracellular targets. The increase in performance when pretraining with peptides underscores the importance of domain-specific datasets when attempting to capture macromolecular interactions. A more stratified investigation of changes made to the model architecture including rotational embeddings, quadratic attention, tokenization strategy, total parameter size, feed-forward dimension, and pretraining data might further elucidate which elements caused the greatest impact to increased prediction.

### Model Design

With the aim of creating a method for encoding noncanonical peptides, we developed a novel tokenization scheme and pretrained a BERT transformer architecture on a database of small molecules and peptides. The resulting model can embed peptides with any modification or non-canonical amino acid by representing macromolecules as Simplified Molecular-Input Line-Entry System (SMILES) strings.

We based our architecture on that used in ChemBERTa^43^ but introduced several changes. We increased the context length of the model from 512 tokens to 768, which allows for encoding of peptides and small proteins up to roughly 100 amino acids in length. Generally, peptide-based pharmaceuticals, such as semaglutide, rarely surpass 50 amino acids in length. We chose to expand this window so that we could encode peptides and small proteins of up to 100 amino acids. This allowed us to take advantage of the SmProt^28^ database for pretraining, as it contained small proteins of up to 100 amino acids. This expanded context window also allowed for training loss to improve more rapidly, as each sequence can contain more tokens and thus a higher number of masked residues per sequence.

We also replaced static sinusoidal positional embeddings with rotational embeddings as developed in RoFormer,^33^ which has been shown to reduce perplexity for protein language modeling in ESM-2.^13^ The vocabulary for tokenization was defined by analyzing n-grams of up to 5 characters to identify common parenthetical insertions and bracketed ions, a method similar to byte-pair encoding^44^ used in natural language modeling. We included the top n-grams of up to four characters. This process generated 581 unique tokens, providing a higher compression rate for peptides compared to single character tokenization used in the ChemBERTa models^26,43^ models. Notably, standard byte-pair-encoding did not improve pretraining for ChemBERTa-2.^43^

### Impact of synthetic data for pretraining

In comparison to the large databases of protein sequences and small molecules, there is significantly less peptide data available for pretraining. The use of synthetic data has been successful for chemical language model pretraining, as in the use of the generated small molecule database ZINC22^45^ for pretraining the BERT-style model MoLFormer.^12^ To supplement the limited amount of available peptide data, we generated 10 million randomized peptide sequences with an updated version of CycloPs.^29^ This number was chosen to create a balance between the two datasets of small molecules and peptides and to increase the number of tokens to approximately 100x the number of model parameters (4.6 billion tokens) with the goal of improving pretraining loss.^46^ The synthetic peptides were of length 1–100 amino acids with a small percentage of noncanonical amino acids, D-conformations, PEGylation, N-methylation, and common cyclizations. All these modifications are used in therapeutic peptides, often for the purpose of increasing stability; N-methylation can increase membrane permeability,^47^ and PEGylation reduces renal clearance, resulting in a longer half-life.^48^

### Clustering and finetuning

In a recent preprint, scaffold-based training/test splits were shown to overestimate the ability of a model to generalize to unseen data. ^49^ In order to best assess our model’s ability to generalize to out-of-distribution peptides, we chose here to use an ML-based clustering approach to avoid data leakage during k-fold analysis. The better performance of PeptideCLM on the *leave-one-cluster-out* testing demonstrates the ability of the model to better generalize. This approach seems to be more informative in laboratory scenarios where the available experimental data may not adequately cover the embedding space of all possible peptides.

### Comparison of architectures and methods

Previous research on language models in biology has primarily focused on small molecules^26,43^ or large proteins,^12,50^ thereby leaving a gap in the modeling of mid-sized peptides. Protein language models have been adapted for use with peptides^14^ but cannot encode modified, non-standard amino acids or cyclizations. Our approach bridges this gap by leveraging the flexibility of SMILES notation and the powerful representation learning capabilities of transformer models.

While the ChemBERTa models are effective for generating representations of small molecules, our results indicate reduced performance when modeling peptides. The ChemBERTa models were selected as benchmarks for comparison with PeptideCLM because they are the only publicly available chemical language models with a sufficiently large input context to encode peptides. ChemBERTa is a chemical language model trained via the self-supervised task of masked language modeling (MLM). In comparison, ChemBERTa-2 has a reduced parameter count and employs linear attention, yet is still able to outperform ChemBERTa. Our study suggests that this improvement arises from an alternative pretraining task of multi-task regression on computationally derived molecular descriptors, as the MLM-trained ChemBERTa-2 underperforms the originial ChemBERTa with our assessment.

An alternative structural deep learning approach by Li, et al.^22^ applied various chemical, 2D, and 3D descriptors to predict PAMPA scores in the Cyclic Peptide Membrane Permeability Database (CycPeptMPDB).^36^ This model achieved similar results to our model when predicting on a random split of the data. We could not make a comparison on our embedding-based data split because the model was not publicly available at the time of this study. This modeling approach used pose estimation, requiring peptide structure predictions. Another recent work explored cyclic peptide conformation changes when crossing a membrane.^51^ Chemical language models are structure agnostic, with relationships learned across the entire molecule during training, which provide an advantage by creating internal representations that mimic pose estimation without requiring structure as input. A systematic analysis predicting membrane penetration for various assays was also reported using standard machine learning methods such as random forest and gradient boosting methods.^23^ This study resulted in an R^2^ of 0.658 for predicting PAMPA values in CycPeptMPDB.^36^

A recent paper^52^ used our pretraining dataset including our synthetic peptides to finetune ChemBERTa-2 for peptides and demonstrated improved transfer learning capabilities for predicting PAMPA scores relative to stock ChemBERTa-2. Note that their strategy yielded an RMSE of around 3, whereas our finetuned models here consistently reach RMSEs of less than 1, regardless of which model was used. This observation suggests that finetuning the entire model, as we did here, performs better than transfer learning on embeddings, even if embeddings have been extracted from a finetuned model.

### Future Directions

Due to using quadratic self-attention, we limited our context window to keep the memory during training manageable for both our pretraining and for an end-user. It may be worth exploring if a larger context window combined with a subquadratic sequence model might be useful. One example of a solution to the issue of memory consumption would be to use a bidirectional state space model^53,54^ to generate embeddings.

The pretraining dataset, while large and diverse, may still not capture the full spectrum of possible peptide modifications found in nature and in synthetic biology. Additionally, the current model architecture, although effective, could benefit from further optimization, such as incorporating pre-layer normalization^55^ to enhance training stability. The application of linear attention^56^ to a model pretrained on peptides is worth investigating to see if it improves training loss.

A further limitation is the reliance on chemical strings for peptide representation. While the Simplified Molecular-Input Line-Entry System (SMILES) provides detailed descriptions of chemical structures, it can be cumbersome for representing large and highly complex peptides. Future work could explore alternative string representations or graph transformer-based approaches, to capture the spatial and functional aspects of peptide structures more effectively. Additionally, availability of datasets with cyclic peptides is limited. However, our model could be applied to linear, canonical peptides as well. We hope to explore this direction in the future.

## 5 Conclusion

PeptideCLM represents a significant advancement in peptide modeling, offering a flexible and powerful tool for predicting diverse properties of peptides. Our method of tokenization and model design is an alternative modeling approach in the field of peptide-based drug discovery. Practical application of language models to noncanonical peptides could be used as an aid to guide drug discovery. The insights gained from exploring alternative modeling approaches to peptides will guide future research and development, expanding the repertoire of computational methods available for therapeutic peptide design.

## 6 Acknowledgments

This work was supported by NIH grant 1R01 AI148419. C.O.W. was also supported by the Blumberg Centennial Professorship in Molecular Evolution and the Reeder Centennial Fellowship in Systematic and Evolutionary Biology at The University of Texas at Austin. Computational analyses were performed using the Biomedical Research Computing Facility at UT Austin, Center for Biomedical Research Support. RRID: SCR 021979. The authors thank Luiz Vieira for support and discussions on the topic of language model pretraining and finetuning.

## 7 Data and Software Availability

Instructions, example scripts, and the PAMPA cluster dataset used in finetuning can be found at https://github.com/AaronFeller/PeptideCLM. Pretraining data has been deposited on Zenodo at https://doi.org/10.5281/zenodo.14194470. Model weights for the full (23M), peptide (11M), and chemical (12M) model have been uploaded to Hugging Face at https://huggingface.co/aaronfeller. The peptide generation library and scripts, with instructions, can be found at https://github.com/AaronFeller/CycloPs_v2.

