## Supplemental Figures and Tables for "Peptide-aware chemical language model successfully predicts membrane diffusion of cyclic peptides"

#### 1 Supplemental Tables

Table S1: Datasets used in PeptideCLM masked language modeling pretraining. Checkmarks denote whether a dataset was used in training the model.

| **Data type (curation source)** | **Count** | **Full Model** | **Peptide Model** | **Small Molecule Model** |
|---|---|---|---|---|
| Generated peptides (CycloPs) | 10M | ✓ | ✓ | |
| Natural peptides (SmProt) | 825K | ✓ | ✓ | |
| Small molecules (PubChem) | 10M | ✓ | | ✓ |
| Patented molecules (SureChEMBL) | 2.2M | ✓ | | ✓ |
| **Total** | **23M** | **23M** | **10.8M** | **12.2M** |

Table S2: Dataset details for CycPeptMPDB used in finetuning. SMILES were generated for all peptides and columns represent the average SMILES length, average permeation score, number of peptides considered permeating, number of peptides considered non-permeating, and the total number of peptides for each step of filtering or each cluster.

| | **Avg. SMILES Length** | **$\mathrm{LogP_{exp}}$ Avg.** | **$\mathrm{LogP_{exp}}$ $\geq -5.5$** | **$\mathrm{LogP_{exp}}$ $< -5.5$** | **Total** |
|---|---|---|---|---|---|
| **CycPeptMPDB** | - | - | - | - | 7,451 |
| **PAMPA only** | 150.7 | -5.87 | 2,854 | 4,087 | 6,941 |
| **Without outliers** | 151.5 | -5.72 | 3,847 | 2,854 | 6,701 |
| **Cluster 1** | 125.8 | -5.57 | 767 | 779 | 1,546 |
| **Cluster 2** | 181.0 | -5.90 | 515 | 950 | 1,465 |
| **Cluster 3** | 116.2 | -6.01 | 402 | 1,053 | 1,455 |
| **Cluster 4** | 134.3 | -5.80 | 413 | 793 | 1,206 |
| **Cluster 5** | 225.3 | -5.06 | 419 | 116 | 535 |
| **Cluster 6** | 210.0 | -5.29 | 338 | 156 | 494 |

Table S3: Hyperparameters for pretraining and finetuning PeptideCLM.

| **Model Architecture** | **Pretraining** | **Finetuning** |
|---|---|---|
| Total Parameters | 44M | 44M |
| Attention Heads | 12 | 12 |
| Number of Layers | 6 | 6 |
| Hidden Dimension | 768 | 768 |
| Feed-forward Dimension | 3,072 | 3,072 |
| Vocabulary size | 586 | 586 |
| Initializer | Xavier | Xavier |
| Optimizer | Adam | Adam |
| $\beta 1$ | 0.9 | 0.98 |
| $\beta 2$ | 0.9 | 0.98 |
| $\epsilon$ | $10^{-8}$ | $10^{-8}$ |
| Learning rate | $5 \times 10^{-5}$ | $5 \times 10^{-6}$ |
| Dropout rate | 0.1 | 0.15 |
| Clip norm. | 0 | 0 |
| Layer norm. $\epsilon$ | $10^{-12}$ | $10^{-12}$ |
| Weight decay | 0 | 0.001 |
| Activation | GELU | GELU |
| Batch size | 64 | 16 |
| Masking | 15% (80/10/10) | - |
| Steps/Epochs | 10 epochs (20 for ablation studies) | 10,000 steps max. |
| Precision | 16-mixed | 16-mixed |
| Data Strategy | Distributed Data Parallel | Single GPU |

### 2 Supplemental Figures

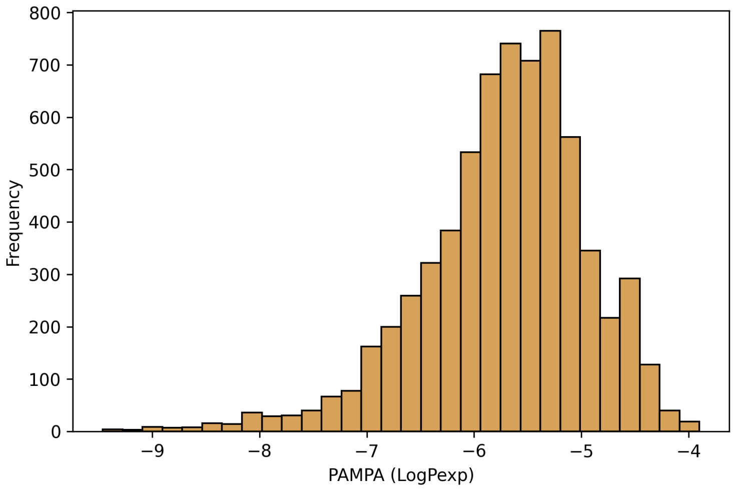


Figure S1: Distribution of Log$P_{exp}$ scores measured by PAMPA assay for cyclic peptides in CycPeptMPDB. Peptides that were undetectable and scored -10 were removed.

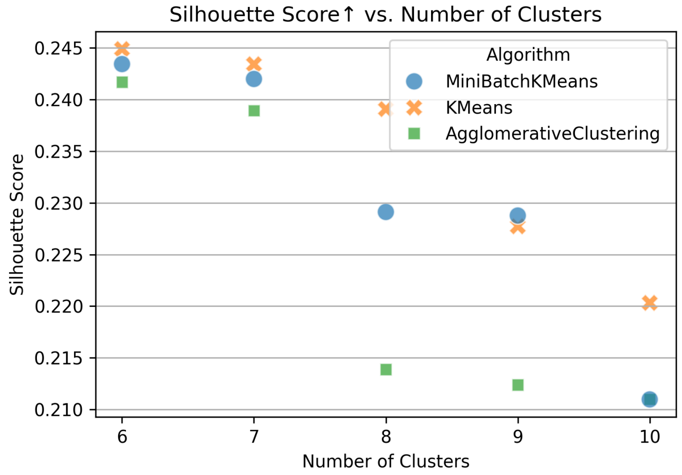


Figure S2: Clustering of PAMPA data was evaluated using Silhouette Score to determine the best clustering method. A higher silhouette score represents higher cohesion and better separation of the clusters. MiniBatchKMeans, KMeans, and Agglomerative Clustering were assessed for sizes of 6 to 10 clusters.

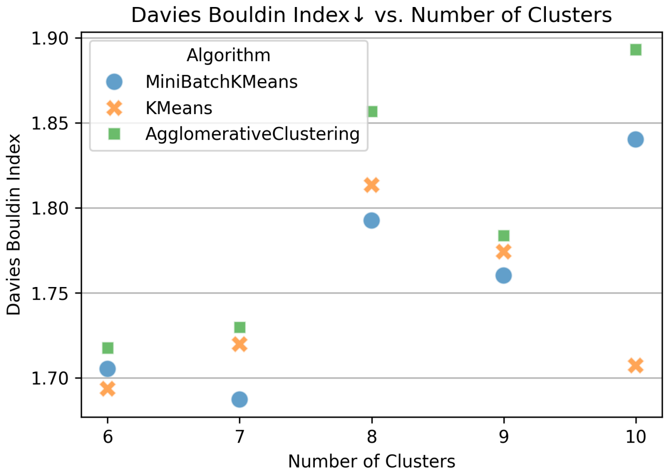


Figure S3: Clustering of PAMPA data was evaluated using Davies-Bouldin Index (DBI) to determine the best clustering method. A lower DBI score shows improved separation and compactness of the clusters. MiniBatchKMeans, KMeans, and Agglomerative Clustering were assessed for sizes of 6 to 10 clusters.

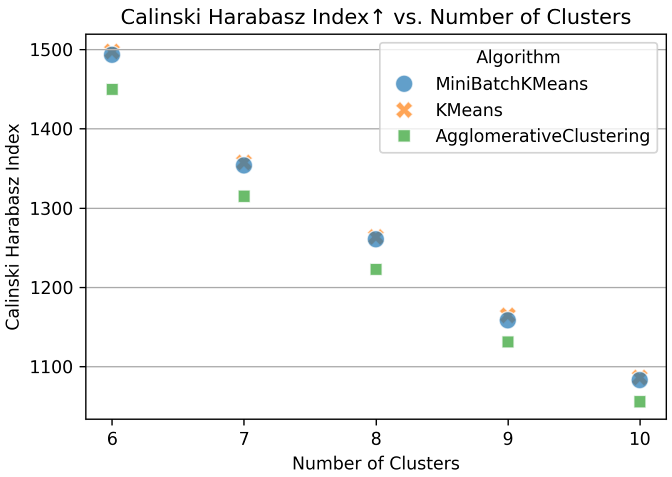


Figure S4: Clustering of PAMPA data was evaluated using Calinski-Harabasz Index (CHI) to determine the best clustering method. A higher CHI score shows improved separation and compactness of the clusters. MiniBatchKMeans, KMeans, and Agglomerative Clustering were assessed for sizes of 6 to 10 clusters.

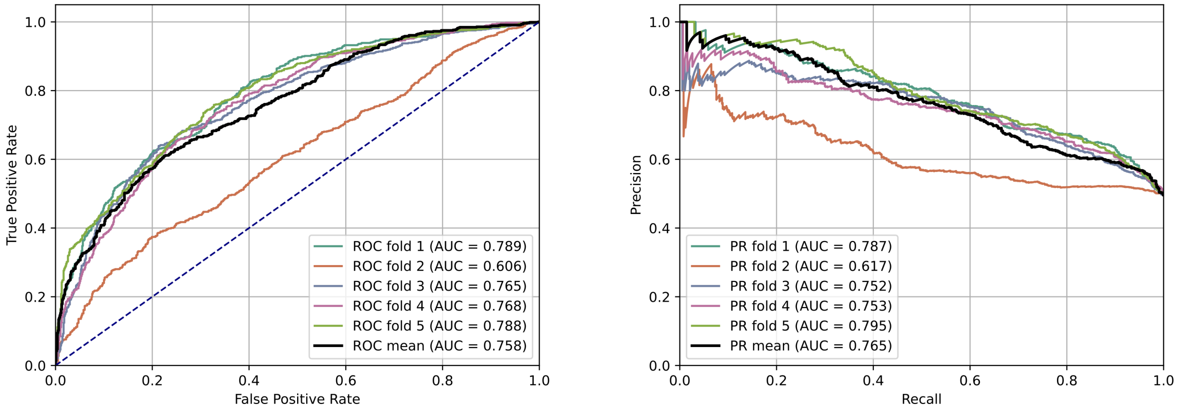


Figure S5: Receiver-operator curves and precision-recall curves for PeptideCLM-23M on holdout test cluster 1.

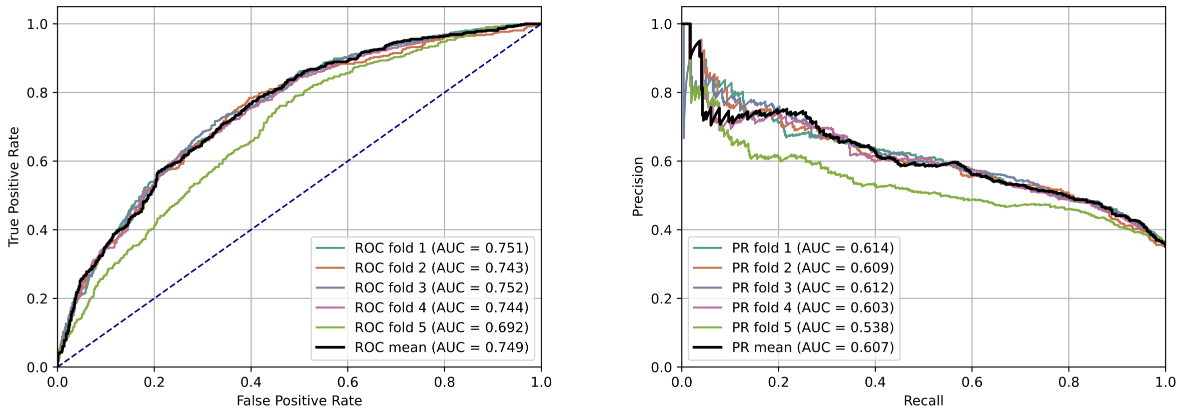


Figure S6: Receiver-operator curves and precision-recall curves for PeptideCLM-23M on holdout test cluster 2.

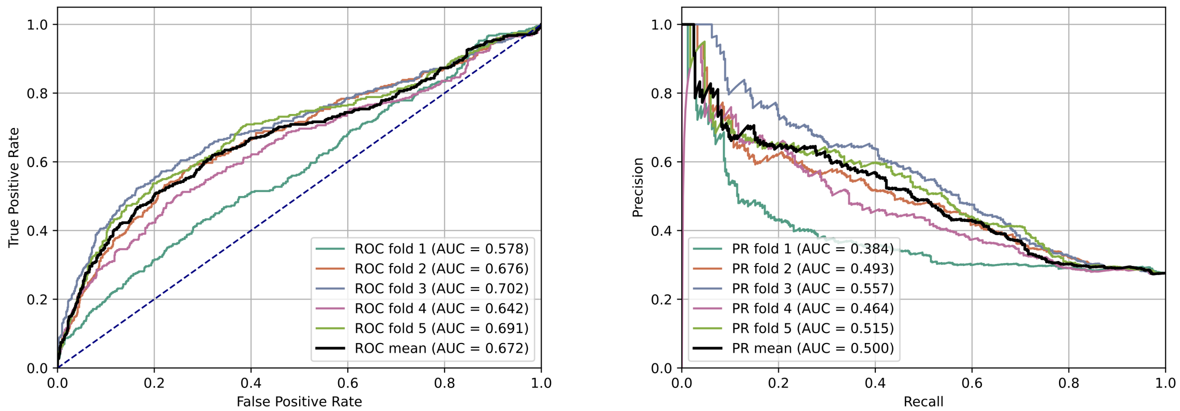


Figure S7: Receiver-operator curves and precision-recall curves for PeptideCLM-23M on holdout test cluster 3.

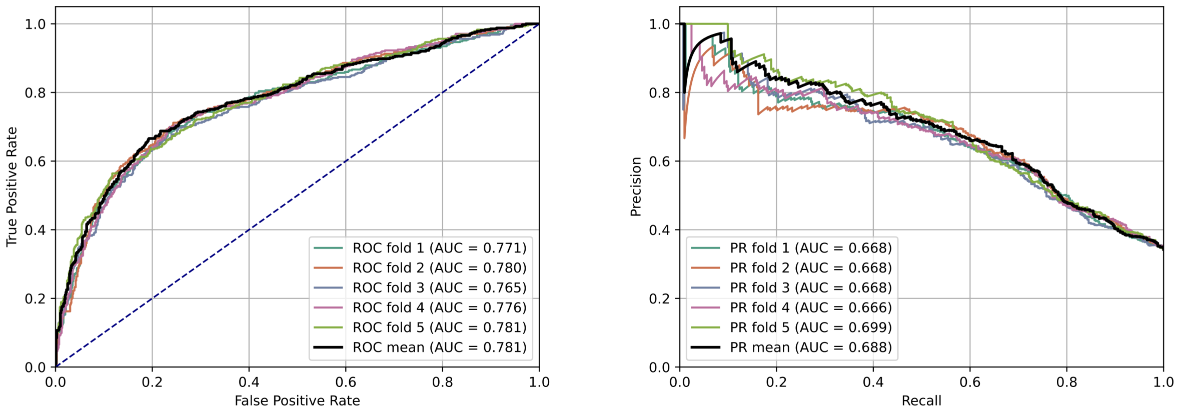


Figure S8: Receiver-operator curves and precision-recall curves for PeptideCLM-23M on holdout test cluster 4.

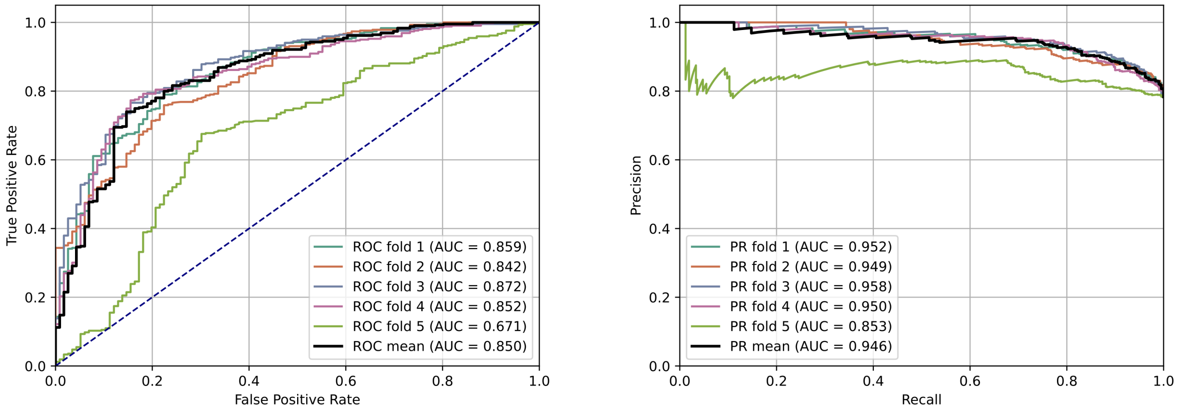


Figure S9: Receiver-operator curves and precision-recall curves for PeptideCLM-23M on holdout test cluster 5.

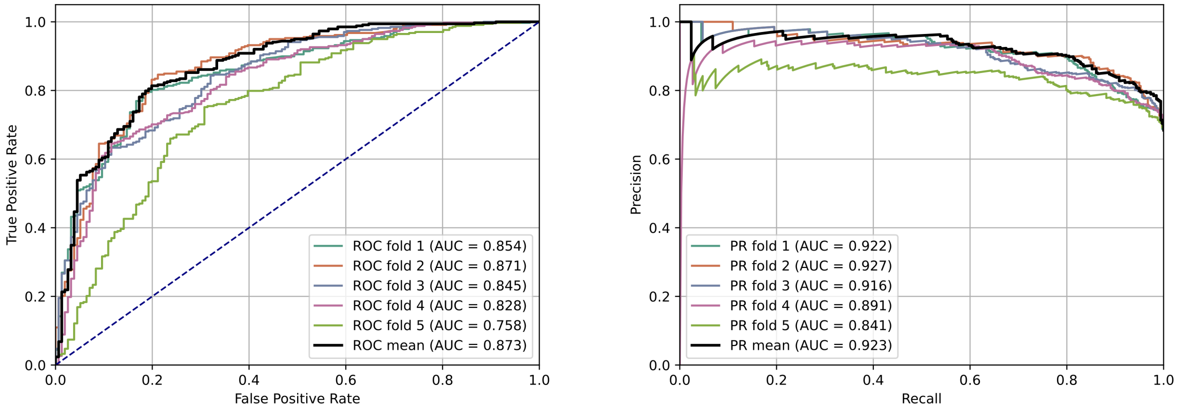


Figure S10: Receiver-operator curves and precision-recall curves for PeptideCLM-23M on holdout test cluster 6.
